# Whole-miRNome sequencing (WMS) - a panel for targeted sequencing of all human miRNA genes

**DOI:** 10.1101/2025.03.09.641621

**Authors:** Paulina Galka-Marciniak, Martyna Olga Urbanek-Trzeciak, Daniel Kuznicki, Natalia Szostak, Adrian Tire, Paulina Maria Nawrocka-Muszynska, Katarzyna Chojnacka, Malwina Suszynska, Katarzyna Klonowska, Karol Czubak, Magdalena Machowska, Anna Philips, Konstantin Maksin, Laura Susok, Michael Sand, Janusz Rys, Jolanta Jura, Magdalena Ratajska, Hanna Dams-Kozlowska, Janusz Kowalewski, Marzena Anna Lewandowska, Piotr Kozlowski

**Affiliations:** Institute of Bioorganic Chemistry, Polish Academy of Sciences, Poznan, Poland; Laboratory of Experimental Medicine, Medical University of Warsaw, Warsaw, Poland; Laboratory of Nuclear Proteins, Faculty of Biotechnology, University of Wroclaw, Wroclaw, Poland; University of Commerce and Services (WSHIU), Medical Department. Poznan, Poland; Department of Dermatology, Venereology and Allergology, Ruhr-University Bochum, Bochum, Germany; Department of Dermatology, Dortmund Hospital gGmbH and Faculty of Health, Witten/Herdecke University, Dortmund, Germany; Department of Dermatology, Venereology and Allergology, St. Josef Hospital, Ruhr-University Bochum, Bochum, Germany; Department of Plastic, Reconstructive and Aesthetic Surgery, St. Josef Hospital, Essen, Germany; Maria Sklodowska-Curie National Research Institute of Oncology, Krakow, Poland; Department of General Biochemistry, Faculty of Biochemistry, Biophysics and Biotechnology, Jagiellonian University, Krakow, Poland; Division of Pathology and Neuropathology, Medical University of Gdansk, Gdansk, Poland; Department of Medical Laboratory Science, University of Otago, Dunedin, New Zealand; Department of Cancer Immunology, Poznan University of Medical Sciences, Poznan, Poland; Department of Diagnostics and Cancer Immunology, Greater Poland Cancer Centre, Poznan, Poland; The Ludwik Rydygier Collegium Medicum, Department of Thoracic Surgery and Tumours, Nicolaus Copernicus University, Bydgoszcz, Poland; The Franciszek Lukaszczyk Oncology Center, Department of Molecular Oncology and Genetics, Bydgoszcz, Poland

**Keywords:** miRNA, somatic mutations, SMAD4, DICER1, miRNA biogenesis, noncoding RNA, miR-142, miRNome, WMS, cancer mutations, miRNA CNA

## Abstract

There is a growing interest in the genetic variation of noncoding genomic elements, including miRNAs, and several mutations in miRNA genes implicated in human diseases, including cancer, have already been detected. However, the lack of dedicated analytical tools severely hampers progress in this area. In this study, we developed whole-miRNome sequencing (WMS), which enables targeted sequencing of all human miRNA genes (n∼2000) and 28 miRNA biogenesis genes. Herein, by sequencing almost 600 samples, including ∼300 tumor/normal pairs of samples from different cancer types, we identified ∼2, 000 mutations, including 1, 435 cancer somatic mutations, with 879 occurring in miRNA genes. These mutations were located in all parts of the genes, including seed or cleavage sites essential for the functioning of miRNA genes. The high reliability of the mutations was confirmed through various approaches, including different sequencing methods. The analysis identified several miRNA genes with functional enrichment of cancer mutations, including *MIR3928*, specifically mutated in basal cell carcinoma (BCC), indicating its potential role in this cancer. WMS also allowed the identification of multiple copy number alterations, hotspots of which often encompassed miRNA genes. WMS provides highly effective, low-cost sequencing of all miRNA genes in different types of samples, including highly degraded FFPE samples.

## INTRODUCTION

miRNAs are short (19-23 nt) single-stranded noncoding RNAs that posttranscriptionally downregulate the expression of the majority of human genes by either translation repression and/or mRNA deadenylation and degradation. Over three decades of research showed that specific miRNAs regulate and control numerous cellular and physiological processes, such as development, cell proliferation, differentiation, and apoptosis, and play a role in the pathogenesis of various diseases (1). The role of miRNAs has been particularly intensively studied in cancer, and many miRNAs playing an essential role in cancer-related processes and consistently either upregulated or downregulated in particular cancer types or specific cancer conditions have been identified (2). Among miRNAs whose role in cancer is best documented are the let-7 family, miR-17-92 cluster (oncomiR-1), and miR-21 (3). Several miRNAs have also been clinically tested as cancer biomarkers, cancer therapeutics, or targets of cancer treatments (4).

miRNAs are generated from long primary precursors (pri-miRNAs) through a multistage process of miRNA biogenesis, of which the most important steps are (i) excision of ∼80 nt long hairpin-shaped part (pre-miRNA) catalyzed by the microprocessor complex which core is composed of DGCR8 and nuclease DROSHA (in nucleus), and (ii) processing of pre-miRNA by the nuclease DICER1 cutting off its terminal loop and generating a miRNA duplex (in cytoplasm). Upon loading into the miRNA-induced silencing complex (miRISC), the miRNA duplex releases one of its strands and uses the other as the guide strand (mature miRNA) that recognizes miRNA targets, usually located in the 3’ untranslated region (UTR) sequence of the mRNA.

miRNA precursors are encoded either within protein-coding genes or by independent transcriptional units, also known as miRNA host genes, that are transcribed by RNA polymerase II. The most crucial and conservative part of miRNA transcriptional units is a ∼100 nt long segment encompassing the hairpin-shaped structure of pre-miRNA, commonly referred to and annotated by the HUGO Gene Nomenclature Committee as the miRNA gene. Many miRNA genes occur in clusters of 2-46 genes in one transcriptional unit (5). There are currently ∼2000 miRNA genes annotated in the human genome, of which ∼600 are well-validated in major miRNA databases, miRBase (6) and MirGeneDB (7).

Despite the enormous interest in noncoding RNA, especially miRNA, very little is known about the genetic variability of miRNA-encoding genes, including common polymorphism, hereditary germline mutations, and cancer somatic mutations in these genes. Nonetheless, a few well-documented cases of mutations in miRNA genes are responsible for Mendelian diseases and are recurrently found in cancer. Examples include mutations in *MIR184* (8–10) and *MIR96* (11, 12), associated, respectively, with hereditary eye diseases and nonsyndromic hearing loss; germline and somatic mutations and deletions of the *MIR15A/MIR16-1* cluster in CLL (13); and somatic mutations in *MIR142* [summarized in (14)] in different hematologic malignancies, particularly in lymphomas (reviewed in (15)). Additionally, there are numerous single-nucleotide polymorphisms (SNPs) located within miRNA genes, some of which (e.g., *MIR125A*, *MIR146A*, *MIR502*, and *MIRLET7* family) were associated with different diseases or disease-related conditions [summarized in (15)]. It was discussed that such variants may affect various aspects of miRNA gene function, including miRNA target recognition (if mutation occurs in the seed sequence), miRNA level, miRNA 5p/3p strand balance, or isomiR predominance (15); however, functional studies mainly were focused on mutations in seed sequences affecting target recognition. Some effects of the mutations are a direct consequence of miRNA sequence changes, but others may result from mutation-induced changes in the structure and stability of miRNA precursors, which affect miRNA biogenesis and processing.

Interest in the noncoding genome has increased with the widespread use of whole-genome sequencing (WGS), but its analysis is still very limited due to the lack of dedicated tools. The limitations concern both computational/statistical methods that allow for identifying, annotating, and interpreting genetic variants in noncoding sequences and experimental techniques, including dedicated sequencing approaches. For example, no such tools are available for noncoding elements, as whole-exome sequencing (WES) is for coding sequences, enabling targeted sequencing of all coding exons in one experiment. The cost of WGS is still high, strongly limiting its applications in terms of the number of analyzed samples and depth of coverage, which is usually relatively low for WGS, hampering some applications.

To meet the limitations of currently used techniques and fulfill the methodological gap for miRNA gene analysis, we developed whole-miRNome sequencing (WMS), a new Next Generation Sequencing (NGS)-based approach enabling targeted sequencing of all miRNA genes complemented by a panel of protein-coding genes playing a role in miRNA biogenesis and functioning (miRNA biogenesis genes). We demonstrated the utility of WMS by sequencing with very high coverage (>700x) several hundred DNA samples of different types and qualities, including high-quality samples extracted from fresh cell cultures, as well as highly degraded, low-quality samples extracted from archival formalin-fixed paraffin-embedded (FFPE) cancer samples. As a result, we detected 2, 016 mutations, including 581 constitutional mutations in cell line samples and 1, 435 cancer somatic mutations. The robustness of the detected mutations was confirmed through various validation methods. 879 (61%) cancer somatic mutations were located in miRNA genes, including highly validated and cancer-related miRNA genes. The mutations were often located in different functional elements of the genes. The analysis showed the over-occurrence and functional enrichment of mutations in some miRNA genes, which may suggest their role in cancer. Finally, we demonstrated the utility of WMS in identifying copy number alterations (CNAs, i.e., deletions or duplications/amplifications. Besides well-known cancer drivers, many identified CNA hotspots also encompassed genes of cancer-related miRNAs.

## METHODS

### Patient sample collection

This study utilized genetic material extracted from the pairs of cancer and the adjacent normal tissue (or, in some cases, the matching blood samples). A total of 286 pairs of tissues were collected at the Franciszek Lukaszczyk Oncology Center, Department of Molecular Oncology and Genetics, Bydgoszcz, Poland [154 FFPE lung cancer adenocarcinoma samples (referred to as LUN), 33 FFPE colon cancer (COL), and 20 ovarian cancer (OVA) samples], at the Medical University of Gdansk, Poland [49 fresh frozen ovarian cancer (OVA) samples with matching control blood samples], at the Center of Oncology, Maria Sklodowska-Curie Memorial Institute, Cracow Branch, Poland [6 fresh frozen renal carcinoma (REN) samples], and at the Department of Plastic Surgery, St. Josef Hospital, Catholic Clinics of the Ruhr Peninsula, Essen, Germany [24 fresh basal cell carcinoma (BCC) tissues placed in RNAlater (Qiagen, Hilden, Germany) and stored at −80°C]. The samples were collected after obtaining signed informed consent from the patients. The study was approved by the Bioethics Committee of the Poznan University of Medical Sciences, Poland (03.08.2018) and was conducted in accordance with the local institutional bioethical committees and the Declaration of Helsinki.

### The WMS panel design

The WMS panel is designed to capture a specific set of targets encompassing 1849 miRNA genes, exons of 28 genes involved in miRNA biogenesis and function, and 10 well-known lung cancer driver genes (or their hotspot-containing exons). Genomic coordinates of all targeted regions are listed in Supplementary Table S1. The WMS probes were designed in the Agilent SureDesign web portal, if possible, utilizing validated probes from SureSelect Human All exome V6+UTR r2 enrichment panel (Agilent Technologies, Santa Clara, CA, USA) or designing new probes according to general SureDesign recommendation. 29 miRNA genes were excluded due to their location in repetitive regions, segmental duplications, and/or low-complexity sequences. For most of the miRNA biogenesis genes, the WMS panel covers the entire coding (CDS), 5’UTR, and 3’UTR sequences. For the following genes 3’UTRs were not covered (*AGO1*, *LIN28A*, *LIN28B*, *SMAD4,* and *TNRC6A*) or were covered partially (*AGO2*, *AGO3*, *SRSF3*). Although probes were designed against the GRCh19 genome assembly, all downstream analyses were performed, and the results are presented against GRCh38. WMS is available from Agilent SureSelect Custom target capture under design ID:3115731.

### NGS and data analysis

Before library preparation, all DNA samples were quantified using a NanoDrop One (Thermo Scientific, Waltham, USA) and Qubit fluorometer 3.0 (Invitrogen Carlsbad, CA, USA) [Qubit dsDNA HS Assay (Life Technologies, Carlsbad, USA)]. The quality/integrity of each sample was assessed with the use of our routine in-lab procedure, based on the image of the sample (50-400 ng) in agarose gel electrophoresis, on a 4-point scale where 3 indicates DNA mainly in the high-molecular-weight (upper band), 2 indicates a smear of different molecular sizes, 1 indicates DNA mostly in the low-molecular-weight fragments (lower band), and 0 indicates no visible signal. Most of the libraries were prepared with 200 ng of DNA using the SureSelectXT Target Enrichment System for Illumina Paired-End Multiplexed Sequencing Library kit (Agilent Technologies, Santa Clara, CA, USA) according to the manufacturer’s recommendations. For a small group of samples (11 pairs), the SureSelectXT High-sensitivity kit was used. Samples were sequenced on Illumina (San Diego, CA, USA), HiSeq 4000 (180 samples) and NovaSeq 6000 (414 samples) with the 100 bp paired-end read mode. Library preparation, enrichment, and targeted sequencing were performed at Macrogen Inc. (Seoul, Republic of Korea). Demultiplexing of the sequencing reads was performed with Illumina bcl2fastq (v 2.20). Low-quality reads were filtered out, and adapters were removed with AdapterRemoval v.1.5.4. The paired-end reads were aligned to the human reference genome (version GRCh38/hg38) using Burrows-Wheeler Aligner (BWA-mem) (16). Duplicated reads were removed, and sequencing metrics were collected with the Picard tools package (v 2.19.0). Indel realignments and base quality score recalibration were performed with GATK version 4.1.2.0. SAM to BAM conversion was performed with SAMtools (v 1.2) (17).

### Samples fingerprinting

To verify the matching of the analyzed pairs of tumor and normal samples, we compared their genotypes based on the analysis of 10 independent (unlinked) SNPs located in distant miRNA and miRNA biogenesis genes (18, 19) and with minor allele frequency (MAF)>0.1 in the general population according to 1000Genomes Project (Supplementary Table S2). Based on the analysis, 7 sample pairs showing signs of mispairing were excluded from further analysis.

### Somatic mutation calling and filtration

Somatic variants were called with MuTect2 version 4.1.0.0. (20) with the default parameters in the tumor-normal mode and with gnomAD resource to filter germline variants based on population allele frequency. We used FilterMutectCalls to filter out variants for which alternative reads’ median base quality and mapping quality were too low (--min-median-base-quality 20 and --min-median-mapping-quality 30). Additionally, to further increase the reliability of the identified mutations (and avoid the identification of uncertain mutations), we filtered out common SNPs (dbSNP release 153) and mutations with variant allele frequency (VAF) <0.05. Finally, we removed variants with SEQQ >20, indicating that alt alleles are not sequencing errors, and GERMQ quality >20, indicating that alternative alleles are not germline variants. The miRNA gene mutations were annotated using a modified miRMut protocol (21) and a set of in-house Python scripts described before (22). A few miRNA genes assigned in miRBase v.22 as ‘dead entries’ were removed from the analysis. Finally, we analyzed 1817 miRNA genes defined as sequences encoding hairpin-shaped pre-miRNAs flanked upstream and downstream by 25 nucleotides as described before (14, 23) and roughly corresponding to/overlapping the miRNA gene designated by the HUGO Gene Nomenclature Committee. All mutations were defined according to HGVS nomenclature at the transcript (for miRNA and protein-coding gene) and/or protein levels (for protein-coding genes), and the effects of mutations in protein-coding genes were predicted using the Ensembl Variant Effect Predictor (VEP) tool (24). Each miRNA gene mutation was assigned with a defined before-weighted functional mutation score (14, 23). The score is based on localization of mutations in miRNA gene functional motifs/subregions, i.e., seed (guide-strand only) (2x); miRNA duplex (1.5x); functional protein-binding motifs, and DROSHA/DICER1 cleavage sites (1.5x); other locations, including flanks, miRNA precursor apical loop (1x).

### Validation of mutations with Sanger sequencing

We selected 43 mutations of different types, different VAFs, and from different tumors to verify the sequencing results. For each of the selected mutations, we designed a pair of primers flanking the putative mutation site and generating the PCR amplicon of ∼400 nt length (Supplementary Table S3). PCR was performed according to the manufacturer’s recommendations (GoTaq DNA polymerase, Promega, Madison, WI, USA). All PCR products were purified using the EPPIC Fast kit (A&A Biotechnology, Gdynia, Poland) and sequenced in two directions using the BigDye v3.1 kit (Applied Biosystems, Foster City, CA, USA). The sequencing reactions were separated with capillary electrophoresis on ABI PRISM 3130xl (Applied Biosystems, Foster City, CA, USA).

### Identification of potential cancer drivers with OncodriveFML

Cancer driver identification was performed with OncodriveFML (v.2.4.0) (25) across all samples (pancancer) and separately for individual cancer types using the CADD score (26). Additionally, the analysis of BCC samples was complemented with the use of the DANN score (27), which should provide enhanced accuracy in predicting the functional consequences of noncoding variants. The signatures were computed by cancer type as a classifier, with the statistical method set to “arithmetic mean”, including indels. The recommended Q-value 0.25 threshold was used for significant results.

### Analysis of Copy Number Alterations (CNAs)

Identification of CNAs in individual samples was performed using CNVkit (28) against reference created from normal samples for each tumor type separately. The segmentation files generated by CNVkit were subsequently analyzed by GISTIC2.0 (29) (with WMS targeted positions indicated as markers) to identify hotspots of deletions and amplifications across the following sets of cancer samples: LUN, OVA, COL, and BCC. In the case of LUN, only samples sequenced with SureSelectXT (n=140) were included; 11 samples sequenced with SureSelectXT High sensitivity kit were excluded. The following parameters were used: threshold for copy number amplifications and deletions, 0.2; confidence level to calculate the region containing a driver, 0.9; broad-level analysis; and the arm-level peel-off method to reduce noise. To improve analysis reliability and reduce false positive signals, we excluded centromeric regions that are highly variable and rich in repetitive sequences. Although, sex assignment has been correctly performed by the used tools, for clarity, the final results are shown for the human autosomal genome.

### Cancer cell line analysis

Cells were grown in appropriate media to a confluency of ∼80%, and DNA was isolated using AllPrep DNA/RNA/miRNA Universal Kit (QIAGEN, Hilden, Germany). WMS was performed as described above using 200 ng of DNA and quality filtering parameters. We reported all sequence variants identified by Mutect2 that had VAF>0.05 and did not overlap with common SNPs (dbSNP release 153).

## RESULTS

### Development of the whole-miRNome sequencing (WMS) panel

In the framework of the study, we designed the first WMS panel. The panel covers 96% of all (1849/1917) and 99% of validated (499/505) miRNA genes annotated in miRBase (22 release) (6), as well as 99% (503/507) of mRNA genes annotated in MirGeneDB (7). The small fraction of miRNA genes not covered by the panel consists mainly of poorly validated genes, often embedded in low-complexity repeat-rich sequences or duplicated genomic regions. Consistently with the HUGO Gene Nomenclature Committee, as miRNA genes, we defined sequences encoding pre-miRNAs (as annotated in miRBase) with their adjacent flanks (∼20 nt). Additionally, we supplemented the panel with 28 miRNA biogenesis genes, including genes playing a role in (i) the transcription of primary miRNA precursors (pri-miRNAs) [*SMAD4*], (ii) pri-miRNA to pre-miRNA processing in the nucleus [*FUS*, *SRSF3* (SRP20), *DROSHA*, *DGCR8*, *DDX5* (P68), *DDX17* (P72), *GSK3B*], (iii) the export of pre-miRNA to the cytoplasm [*XPO5* (EXP5), *RAN*], (iv) pre-miRNA processing and miRNA maturation [*DICER1* (DICER), *TARBP2* (TRBP), *PRKRA* (PACT), *ADAR*, *KHSRP* (FUBP2, KSRP), *LIN28A*, *LIN28B*, *ZCCHC11* (TUT4), *ZCCHC6* (TUT7), *DIS3L2*], and (v) miRNA:target recognition/interaction and execution of downstream silencing effect [*AGO1*, *AGO2*, *AGO3*, *AGO4*, *GEMIN4*, *MOV10*, *FMR1*, *TNRC6A* (GW182)] [ as in (22)]. Finally, we included in the panel 10 well-known cancer-related genes (or their hotspot-containing exons) listed as most frequently mutated in lung cancer (*BRAF, EGFR, ERBB2, HRAS, KRAS, MAP2K1, MET, NF1, NRAS, RIT1*)(30) to serve as an internal control of highly mutated regions (Figure 1A). In total, the WMS panel encompasses 499, 590 bp covered/baited by 16, 302 probes (Agilent Tier 1), including 333, 174 bp (11, 778 probes) of miRNA genes and 166, 416 bp (4, 524 probes) of miRNA biogenesis and cancer gene exons (Figure 1B).

**Figure 1.**
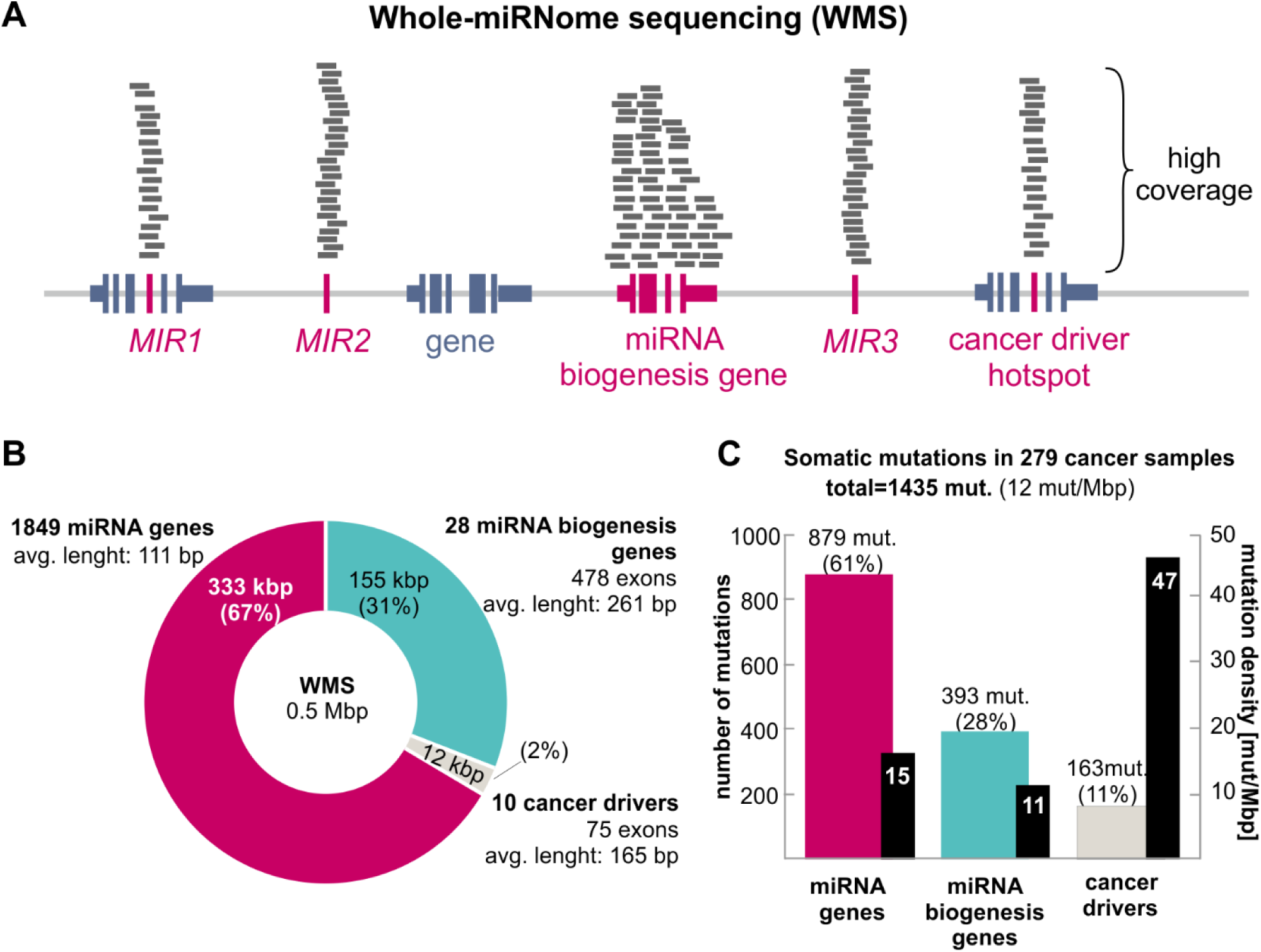
WMS panel design and performance. A) Schematic representation of regions (marked in magenta) enriched by WMS. B) The proportion and number of regions targeted by WMS. C) The number (color bars, left-axis) and density (black bars, right-axis) of somatic mutations identified with WMS in 279 cancer samples, in miRNA genes, miRNA biogenesis genes, and cancer driver regions; mut/Mbp an average number of mutations per million nucleotides per sample.

### WMS performance

To test the panel performance, we run WMS on a total of 580 samples: (i) 279 (tumor/normal) pairs of cancer samples, including 151 lung adenocarcinomas (LUN), 68 ovarian carcinomas (OVA), 31 colorectal cancers (COL), 23 skin basal cell carcinomas (BCC), and 6 renal carcinomas (REN) and (ii) 22 established cancer cell lines. Most (n=202) of the cancer sample pairs were extracted from FFPE blocks, but some (n=77) were extracted from fresh (RNAlater-preserved) or fresh frozen tissues (fresh tissue, FT). DNA from cancer cell lines (n=22) was extracted directly from collected cells. As FFPE samples are inherently of low quality at various stages of degradation, therefore before analysis, the integrity of DNA samples was briefly determined with our internal lab-established procedure on a 4-point scale (from 3 - highest quality to 1 - lowest quality, and 0 - not detectable on agarose gel). The matching of tumor/normal sample pairs was confirmed by SNPs fingerprinting (see Methods).

With some minor exceptions, libraries were prepared with 150-200 ng of DNA and were sequenced to obtain 30 million (M) total reads per sample (3 Gbp). The essential characteristics of WMS performance, i.e., target coverage, duplication rate, target fold enrichment, and number of detected somatic mutations, are shown in Figure 2 and Supplementary Table S4. The mean sample coverage of all samples averaged 732x (range 60-1906x) and, as expected, was somewhat higher for high-quality samples extracted from cell lines (average 1644x) than for archival fresh and fresh frozen tissues and FFPE-extracted samples (Figure 2A). Also, FFPE samples with higher integrity scores showed higher coverage than those with lower ones (Figure 2A). The differences in the coverages may be at least partially explained by the duplication rate, which, as shown in Figure 2A (compare the first two panels) and Figure 2B, is lower in high-quality samples and roughly inversely correlated with coverage. Despite the differences, the coverage of the vast majority (90%) of samples, including FFPE-extracted samples, was >200x, which is much higher than that of standard WES or WGS experiments. Even though the miRNome panel is relatively small ∼0.5 Mbp (∼6.200x smaller than the human genome), the baited region, on average, has been enriched (over a genomic background) by 2248x, and the enrichment did not differ substantially between the sample types (Figure 2A), indicating the robustness of WMS.

**Figure 2.**
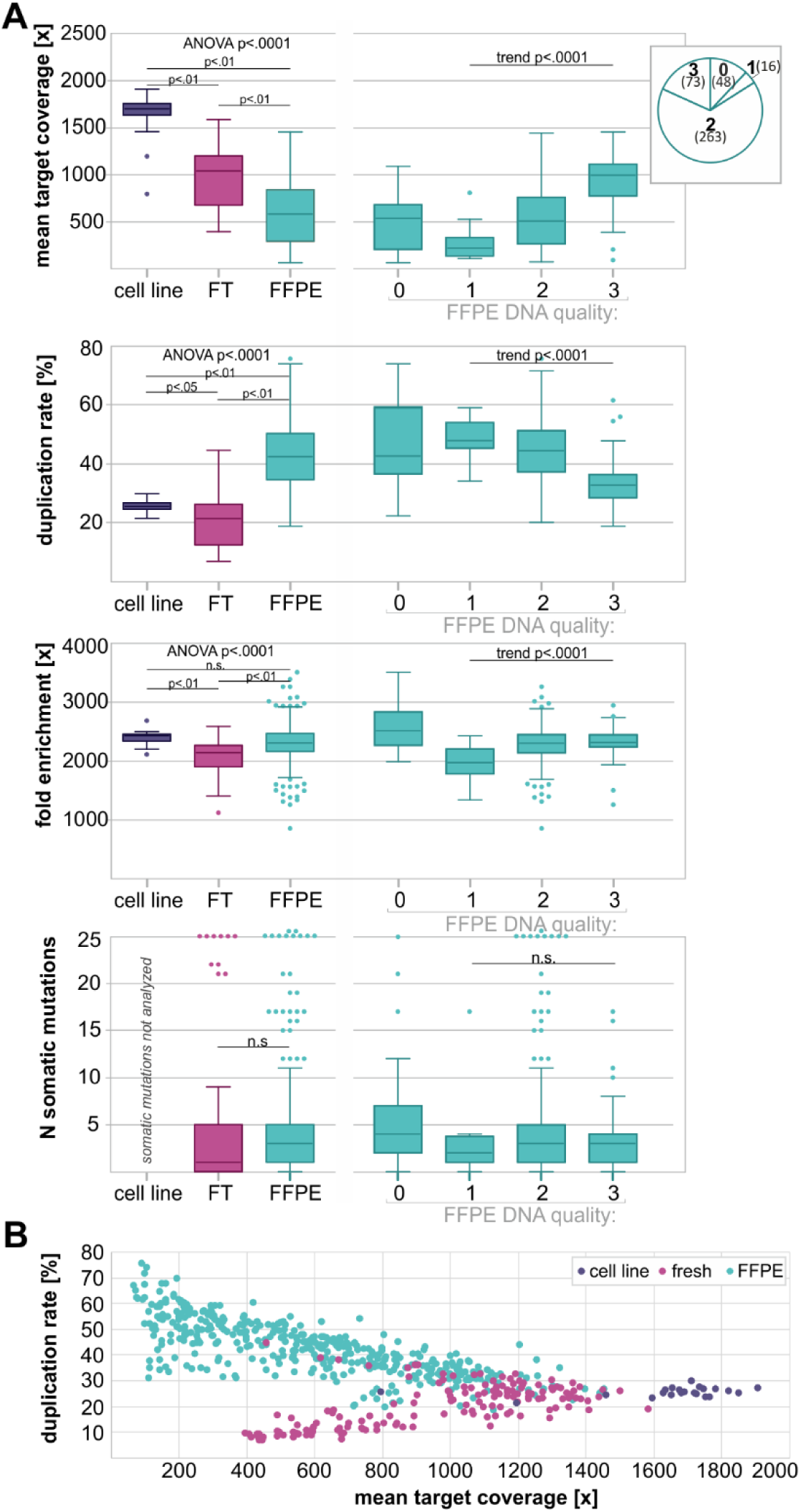
Sequencing metrics and performance of WMS. A) On the left, Tukey’s box and whisker plots showing distributions of mean sequencing coverage, fold enrichment, duplication rate, and number of somatic mutations in DNA samples extracted from cell line, freshly preserved tissue (FT), and FFPE preserved samples; the global ANOVA p-values and post-test p-values for particular comparisons are indicated above the charts. On the right, FFPE samples were additionally divided into 4 DNA integrity/quality categories (see Methods); the ANOVA for trend p-values for categories 1 to 3 are indicated above the graphs; proportions of FFPE samples in the particular integrity categories are shown in inset (pie-chart). B) A scatter plot shows the relationship between mean sequencing coverage and duplication rate for different types of samples analyzed.

We called somatic mutations with the Mutect2 algorithm and processed them with the miRMut protocol(21). In total, in 279 cancer samples, we identified 1, 435 somatic mutations, of which 879 (61%) were located in miRNA genes, 393 (28%) were located in miRNA biogenesis genes, and 163 (11%) in the control cancer-driver genes (Figure 1C, Table 1 and Supplementary Table S5). Despite some dependence of WMS performance parameters on DNA quality (above), we did not observe any effect of DNA quality on the number of detected mutations (Figure 2A).

**Table 1.** Summary of mutations identified in analyzed cancer types.

|  | All samples | LUN | OVA | COL | BCC | REN |
| --- | --- | --- | --- | --- | --- | --- |
| **N pairs** | 279 | 151 | 68 | 31 | 23 | 6 |
| **total N of mut.** | 1435 | 715 | 277 | 240 | 199 | 4 |
| **N (%) mutated samples** | 227 (81) | 133 (88) | 43 (63) | 26 (84) | 22 (96) | 3 (50) |
| **N (%) substitutions** | 1312 (91) | 651 (91) | 256 (92) | 214 (89) | 188 (94) | 3(75) |
| **N miRNA mut.** | **879** | **426** | **164** | **141** | **145** | **3** |
| **N (%) mutated samples** | **187 (67)** | **116 (77)** | **30 (44)** | **16 (52)** | **22 (96)** | **3 (50)** |
| **N (%) substitutions** | **825 (94)** | **395 (93)** | **155 (95)** | **133 (94)** | **140 (97)** | **2 (67)** |
| **N biogenesis mut.** | **393** | **184** | **85** | **72** | **51** | **1** |
| **N (%) mutated samples** | **120 (43)** | **66 (44)** | **25 (37)** | **12 (39)** | **16 (70)** | **1 (17)** |
| **N (%) substitutions** | **348 (89)** | **169 (92)** | **74 (87)** | **54 (75)** | **50 (98)** | **1 (100)** |
| **N cancer mut.** | **163** | **105** | **28** | **27** | **3** | **-** |
| **N (%) mutated samples** | **116 (42)** | **83 (54)** | **13(19)** | **18 (58)** | **2 (9)** | **-** |
| **Sample type** | - | FFPE | FFPE, FT | FFPE | FT | FT |

### WMS validation

First, to estimate the fraction of false-positive mutations, we resequenced (with Sanger sequencing) 43 mutations in both miRNA and protein-coding genes representing different mutation types. The analysis confirmed 42 mutations, indicating a very low (2%) fraction of potential false-positive mutations. Next, we compared the frequency of mutations in well-established lung adenocarcinoma driver genes in our LUN samples and the previous large-scale study of lung adenocarcinoma reported by TCGA (30). The analysis showed 32% of samples with mutations in *KRAS*, 14% in *NF1*, 8% in *EGFR*, 3% in *BRAF*, 1% each in *ERBB2*, *MET*, and *NRAS*, and no mutations in *MAP2K1*, which is consistent with the TCGA study (30) (r2=0.932; p < 0.0001) Supplementary Figure S1. Moreover, as *EGFR* hotspot driver mutations are biomarkers in cancer treatment and are routinely tested in lung cancer samples, we compared our results with results of clinical testing performed before on the same samples. As shown in Supplementary Table S6, WMS detected all 12 mutations detected before with the use of commercial high-sensitivity RT-qPCR assays (6x exon 19 in-frame deletions, 3x exon 21 L858R, 2x exon 21 L861Q and exon 19 S768I). Additionally, with WMS, we detected one exon 19 in-frame deletion (L747_S752del) missed in clinical testing. Finally, for 23 BCC samples, we compared the current WMS results with our previous results (31), in which the same samples were sequenced with the use of the commercial WES (SureSelectXT Human All Exon V6 kit (Agilent)). These two data sets substantially differ in sequencing depth, which in the compared region covered by both assays (28 miRNA biogenesis genes) was 701x and 183x for WMS and WES, respectively. Despite this difference, the results of both WMS and WES were very consistent; WMS identified all 37 mutations detected by WES and additionally detected 2 mutations not detected before. The new mutations could most likely be detected due to much higher coverage in WMS.

### General characteristics of the somatic mutations identified in miRNA genes

All (n=879) mutations identified in miRNA genes were annotated and characterized using miRMut, a specialized pipeline dedicated to accurate annotation of genetic variants in miRNA genes developed previously in our lab (21). The mutations were identified in 635 miRNA genes, of which 242 are considered high confidence according to miRBase and/or miRGeneDB, and 53 were classified as cancer-related in the Cancer miRNA Census (CMC (2)) (CMC-miRNA). At least one miRNA gene mutation was detected in 187 (67%) cancer samples, 63 (34%) samples had more than one mutation, and 10 (5%) had more than 10 mutations. The vast majority (93.9%) of identified mutations were substitutions, 3.2% were deletions, 1.5% were insertions, and 1.5% were dinucleotide substitutions (DNS). The frequency and type of mutations in different cancer types are summarized in Table 1.

To look closer at the identified mutations’ localization, we categorized them into different miRNA precursor subregions (Figure 3A) and superimposed them onto the consensus miRNA precursor structure (Figure 3B). As shown in Figure 3A, the mutations are more or less evenly distributed along the entire miRNA precursor sequence (average density 15.3 mut/Mbp). There is no evidence (binomial distribution) of unequal distribution of mutations between the functional miRNA gene subregions (range from 13-18 mut/Mbp). Of the identified mutations, 185 (21%) were located in mature miRNAs (guide strands), with 56 of them being in the seeds, which are the most crucial part of the miRNA gene. It was shown before that seed mutations severely affect miRNA:target recognition [examples in (15)]. As shown in Figure 3D for 5 exemplary miRNA seed mutations identified in CMC-miRNAs (miR-1-3p, miR-101-3p, miR-139-5p, miR-146b-5p, and miR-34c-5p), the seed mutations severely affect the spectrum of predicted target genes, which is consistent with our previous results (14, 23). Another 66 mutations were located in DROSHA and DICER1 cleavage sites, including 47 mutations altering GYM, a motif essential for efficient DICER1 cleavage (32). The prominent examples of such mutations are n.45G>A in *MIR30D* and n.41G>T in *MIR320A*, which dramatically lower the GYM score from 73 to 18 and from 58 to 11, respectively, thus are likely to decrease the precursor processing efficiency. Additionally, 40 mutations were located in sequence motifs recognized by regulatory RNA-binding proteins, which may affect the accuracy and efficiency of miRNA precursor processing (Supplementary Table S5).

**Figure 3.**
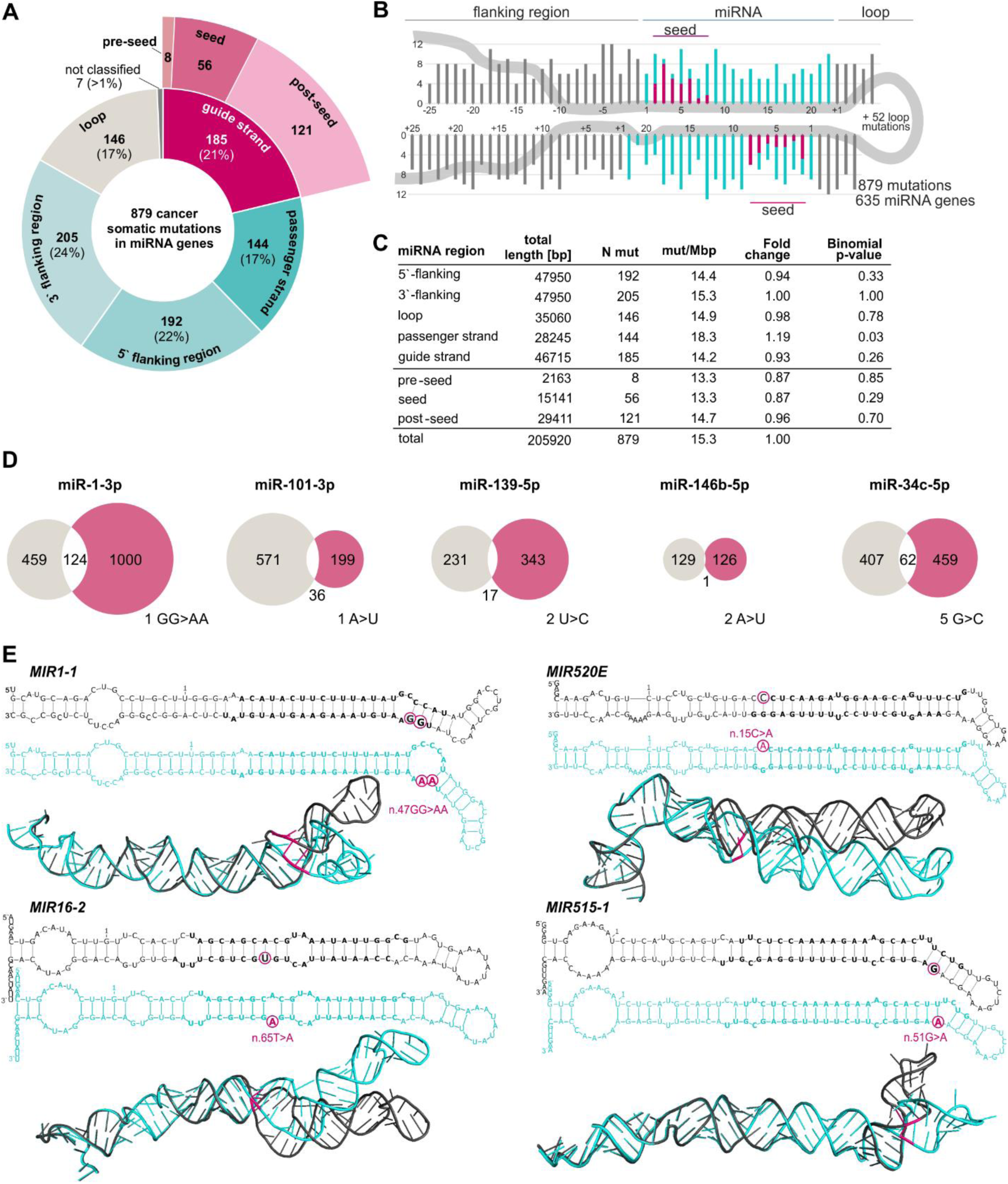
Cancer somatic mutations identified in miRNA genes. A) Distribution of the mutations among miRNA precursor subregions. B) The distribution of the identified somatic mutations along the consensus miRNA precursor sequence. miRNA duplex positions are indicated in turquoise, seed regions in magenta, flanking regions, and the apical loop in gray. If present, mutations localized beyond position 22 in mature miRNAs are shown cumulatively at position 22. As the size and structure of apical loops differ substantially among miRNA precursors, the plot shows only the loop’s first and last five positions; the number of remaining mutations is indicated within the loop. C) Summary and statistics of the mutations identified in subregions of miRNA precursors. D) Venn diagrams showing the effect of representative mutations located in miRNA seeds on the number of predicted (TargetScan 5.2 Custom) targets, as examples were selected miRNAs annotated in CMC(2). The gray and magenta circles indicate the targets predicted for the wild-type and mutant seeds, respectively. The mutated seed positions and the nucleotide changes are shown next to the diagrams. E) Exemplary mutations affecting the secondary 2D (above) and spatial 3D (below) structures of miRNA precursors. Wild-type and mutant structures are shown in black and turquoise, respectively; miRNA duplexes are indicated in bold; mutation positions are indicated in magenta.

In addition to affecting the sequence of functional elements and regardless of location, most miRNA gene mutations also affect the structure and stability of miRNA precursors. Examples of such mutations are shown in Figure 3E.

### Identification of recurrently mutated and potential cancer-driver miRNA genes

To demonstrate the usefulness of WMS, first, we performed the identification of overmutated miRNA genes. The analysis showed that 5 miRNA genes were significantly overmutated (binomial distribution p<0.0003), with ≥ 5 mutations or ≥ 5.5 functionally-weighted miRMut mutation score. Among them were *MIR548F4* with 7 mutations (miRMut score 7.5), *MIR6503* with 5 mutations (miRMut score 5.5), *MIR4445* with 4 mutations (miRMut score 5.5), *MIR4787* with 4 mutations (miRMut score 6.5), and *MIR520E* with 4 mutations (miRMut score 6) (Figure 4A and Supplementary Table S7).

**Figure 4.**
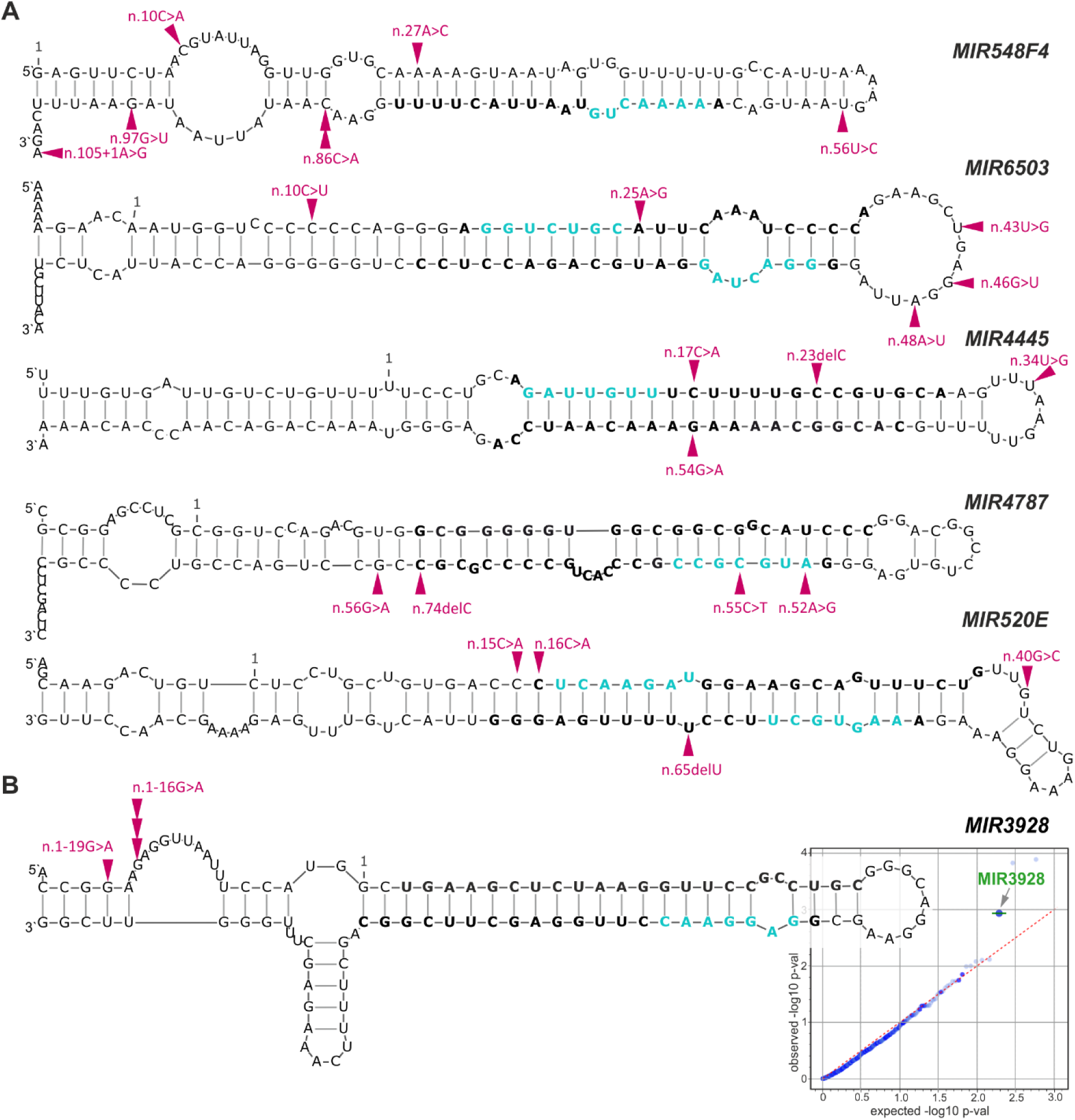
Recurrently mutated miRNA genes. A) Localization of mutations in miRNA genes with 5 or more cancer somatic mutations or with 5.5 or higher functionally-weighted mutation scores. The structures were modeled with Mfold for reconstructed pre-miRNAs extended by 25 nt upstream and downstream flanking sequences. Magenta arrowheads indicate the mutation positions; bold fonts indicate miRNA duplexes; and blue fonts indicate seeds of mature miRNAs. The first nucleotides of miRNA precursors annotated in miRBase and referred to in HGVS nomenclature are indicated as position 1. B) The position of mutations in *MIR3928* gene which was identified as a potential cancer driver. All mutations were identified in BCC. The quantile-quantile (QQ) plot showing the result of OncodriveFML analysis (with the CADD score) visualized as the distribution of expected (x-axis) and observed (y-axis) p-values corresponding to functional mutation enrichment in particular miRNA genes, based on mutations detected in all cancer samples (pancancer analysis). The green color indicates genes defined as significant (q<0.25), according to the OncodriveFML recommendation.

Next, we analyzed the mutations identified in miRNA genes with OncodriveFML, a tool for identifying potential cancer drivers (25). OncodriveFML takes advantage of the functional mutation scores (e.g., CADD, a tool for scoring the deleteriousness of mutations (26)) for the identification of functional mutation biases (enrichment over random chance) in tested regions. OncodriveFML is recommended for both coding and noncoding regions. The cumulative analysis across all cancer samples (pancancer analysis) identified one candidate driver miRNA gene, *MIR3928* (Figure 4B). Further analysis of individual cancer types showed that the cancer driver potential of *MIR3928* is specific to BCC, a cancer type in which *MIR3928* is mutated in 4 (out of 23) samples (Supplementary Figure S2A). The driver potential of *MIR3928* in BCC was also confirmed using DANN (Supplementary Figure S2B), an alternative score for annotating the pathogenicity of genetic variants with improved performance in noncoding regions (27). *MIR3928* was mutated with 4 mutations (miRMut score 4) in 4 BCC samples. The mutations were located in two nearby positions at the 5’ flanking region, n.1-16G>A and n.1-19G>A (Figure 4A). Three of the mutations (with VAF >20%) were analyzed and confirmed by Sanger sequencing. Other miRNA genes identified as potential cancer drivers in individual cancer types include *MIR548AG2* in BCC (in total 4 mutations, miRMut score 5), *MIR3138* in LUN (in total 3 mutations, miRMut score 3.5), and *MIR507* (in OVA (in total 3 mutations, miRMut score 3) (Supplementary Figure S2).

### Identification of mutations in miRNA biogenesis genes and potential cancer drivers

In the panel of 28 miRNA biogenesis genes, we identified 393 mutations, including 30 (7%) in 5’UTRs, 234 (60%) in CDSs, and 129 (33%) in 3’UTRs. Among the CDS mutations were 147 missense mutations, 52 synonymous mutations, and 32 definitive deleterious mutations, including 11 frameshift, 10 nonsense, and 11 splice-site mutations (Figure 5A and Supplementary Table S5). At least one mutation was detected in 120 (43%) samples. The frequency of mutated samples ranged from 17% in REN samples to 70% in BCC samples (Table 1), roughly corresponding to the mutational burden in the analyzed cancer types. Among the most highly mutated genes were *DICER1* with 32 mutations (of which 11 were localized in CDS), *FMR1* with 30 mutations (16 in CDS), *ADAR* with 28 mutations (14 in CDS), *DROSHA* with 27 mutations (22 in CDS), and *SMAD4* with 15 mutations (11 in CDS)(Figure 5B and Supplementary Table S5).

**Figure 5.**
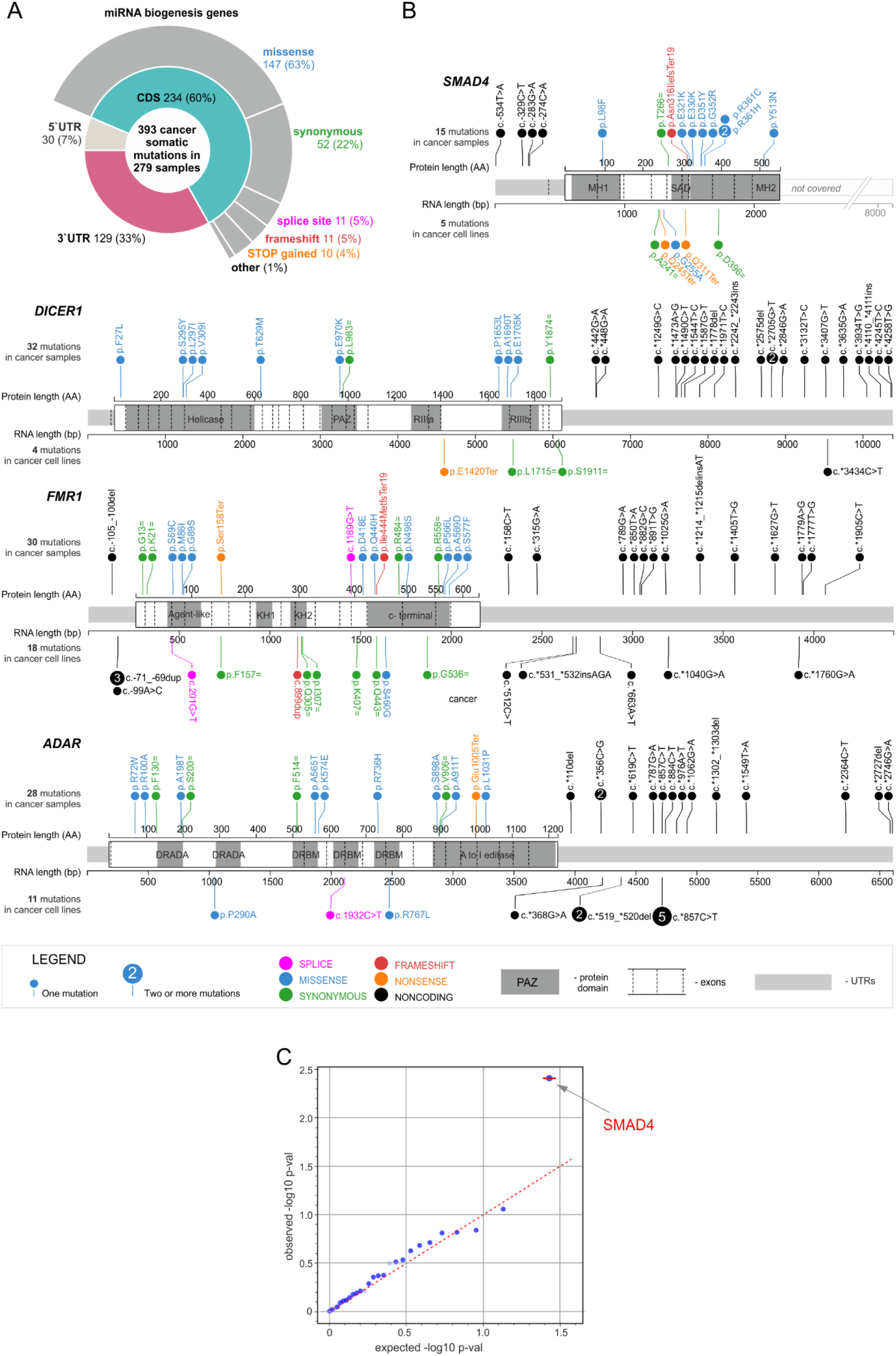
The summary of somatic mutations detected in miRNA biogenesis genes. A) A doughnut chart showing types and subregions of mutations in miRNA biogenesis genes. B) Maps of the most frequently mutated miRNA biogenesis genes. The cancer somatic mutations and constitutional mutations detected in cancer cell lines are indicated as lollipops above and below the maps, respectively; the mutation types are indicated in colors (see the legend). C) The QQ plot showing the result of OncodriveFML analysis (with the CADD score) visualized as the distribution of expected (x-axis) and observed (y-axis) p-values corresponding to the functional enrichment bias of coding mutations in miRNA biogenesis genes, based on mutations detected in all cancer samples (pancancer analysis). The red color indicates genes defined as highly significant (q<0.1), according to the OncodriveFML recommendation.

The analysis of the CDS mutations using Oncodrive FML (Figure 5C) showed a very strong and distinct signal for *SMAD4* as a potential cancer driver (q<0.1). This result further confirms the reliability of mutations detected by the WMS panel, as *SMAD4* is a very well-known tumor suppressor gene playing a role and being frequently mutated in the pancreas, colon, and other intestinal duct cancers, as well as in lung adenocarcinoma (22, 33, 34). Consistently with previous results, most *SMAD4* mutations identified in our study were found in LUN (8 CDS mutations) and COL (4 CDS mutations), and most of them clustered in the MH2 domain, including well-recognized hotspot mutations p.R361C, p.R361H, and p.G352R known to disrupt SMAD4:SMAD2 heterotrimer formation, consequently affecting SMAD4 transcription factor activity (22, 35–37). Another known cancer driver mutation identified in our study is p.E1705K in *DICER1* found in the COL sample. The mutation was observed before in various cancers, predominantly in uterine/endometrial cancer(38, 39). The p.E1705K mutation is located in the catalytic center of the RNase IIIb (RIIIB) domain at the metal ion-binding residue, and it was shown to affect the proper processing of miRNA precursors (22, 40, 41).

### Copy Number Alteration (CNA) analysis

Despite the low number of targeted regions (n ∼ 2000) and a limited number of sequenced cancer samples, we attempted to use WMS for chromosome-arm-level CNA analysis and identification of hotspot deletions and amplifications. As shown in Figure 6A for LUN and in Supplementary Figure S3 for COL, OVA, and BCC, the analysis allowed the detection of numerous somatic CNAs in cancer samples. The analysis also correctly predicted (based on the copy number of sex chromosomes) the gender of the analyzed samples. As expected, CNAs did not occur or were very rare in normal samples (Figure 6A and corresponding panels in Supplementary Figure S3). Since normal samples were mainly extracted from the same tissue types as the corresponding cancer samples and were treated the same way, the level of CNA observed in the normal samples may roughly reflect the level of false positive results. The patterns of recurrent/hotspot CNAs were generally consistent with those observed before for corresponding cancers (30, 42, 43); (Figure 6B and Supplementary Figure S3). For example, we identified 22 LUN hotspot deletions and 27 amplifications (Figure 6B). Among the hotspot amplifications were the regions encompassing *EGFR* (ch7p11)*, KRAS* (ch12p12), and *MCL1* (ch1q21), well-recognized cancer drivers, playing a crucial role and known for being amplified in lung adenocarcinoma (30). Many of the identified CNAs also included well-known cancer-related miRNA genes, including *MIR92B* (ch1q21), *MIR30B* (ch8q24), and *MIR30D* (ch8q24), whose frequent amplifications in lung adenocarcinoma were reported before (44) and many others not reported before (Figure 6B). Among the miRNA genes located in hotspot deletions are *MIR140*, *MIR132*, *MIR212, and MIR22*, playing the role of tumor suppressors (45–47), and the entire chromosome-19 cluster (C19MC) consisting of 46 miRNA genes involved in the regulation of cell cycle and proliferation (48), including *MIR520E*, which we identified as frequently mutated in the LUN samples (Figure 4A). The results of the CNA analysis for other cancer types are shown in Supplementary Figure S3. Of note, in the BCC samples, consistently with our previous study (31) performed using WES, we identified the recurrent deletion of chromosome 9q and the recurrent duplication/amplification of chromosome 9p. Chromosome 9q includes tumor suppressor *PTCH1*, the most crucial driver gene for BCC, and a cluster of 3 miRNA genes (*MIRLET7A*, *MIRLET7B*, and *MIRLET7F*), members of the tumor suppressor let-7 family (49) while chromosome 9p includes *CD274* (encoding PD-L1), *CD273* (encoding PD-L2), and *JAK2* oncogenes. All chromosome 9 alterations in individual samples were consistent with our previous study, in which they were independently confirmed with MLPA(31). Detailed lists of all identified CNAs are shown in Supplementary Table S8.

**Figure 6.**
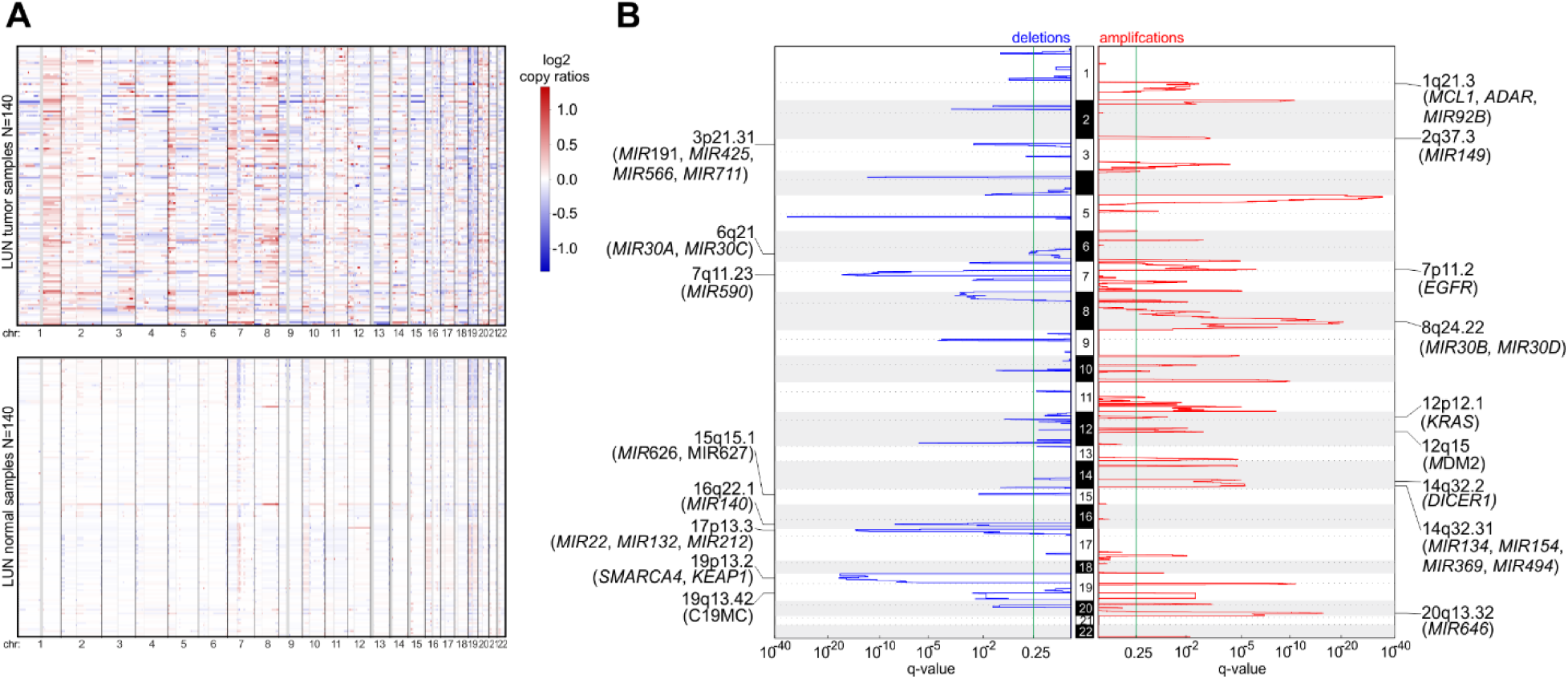
CNA analysis with the use of WMS data in LUN samples. A) A heatmap displaying individual CNAs in tumor (top) and normal (bottom) samples; blue and red indicate deletions and duplications/amplifications, respectively; color intensity indicates log2 of copy-number amplitude. Each row represents an individual sample. B) A plot indicating chromosomal locations (vertical axis) of recurrent amplification (red) and deletion (blue) peaks detected by GISTIC2.0. The height of the peaks (horizontal axes) indicates the significance (q-value) of the region amplification/deletion recurrence. The green line indicates the recommended significance threshold, q=0.25. The selected deleted and amplified regions/genes are indicated on the graphs.

### WMS of cancer cell lines

Cell lines are frequently utilized for the functional characterization of various miRNAs, and miRNA gene mutations can impact these analyses. Therefore, as an additional example of a WMS application, we characterized 22 established cancer cell lines (listed in Table 2). In total, we identified 412 mutations in miRNA genes (75% of which were unique), 138 mutations in miRNA biogenesis genes, and 31 in the control cancer-driver genes (the distribution of mutations in the most frequently mutated genes is shown in Figure 5B). The number of miRNA gene mutations ranged from 91 in SCC-25 (tongue squamous cell carcinoma) to 4 in the SKNMC (bone metastatic neuroblastoma). Among the miRNA genes mutated in the cell lines, 50 were annotated in CMC, including, *MIR21*, *MIR146B*, *MIR150*, and *MIR345,* with mutations in the miRNA duplex; *MIR142*, *MIR200C*, and *MIR335*, with mutations in flanking sequences; and *MIR101-1*, *MIR26A1*, *MIR30A*, *MIR106B*, *MIR221*, *MIR152*, *MIR224*, and *MIR27A* with mutations in apical loops of miRNA precursors. The list of all mutations in miRNA and miRNA biogenesis genes (Supplementary Table S9) may serve as a resource for selecting an appropriate cell line for the functional analysis of specific miRNAs.

**Table 2.**
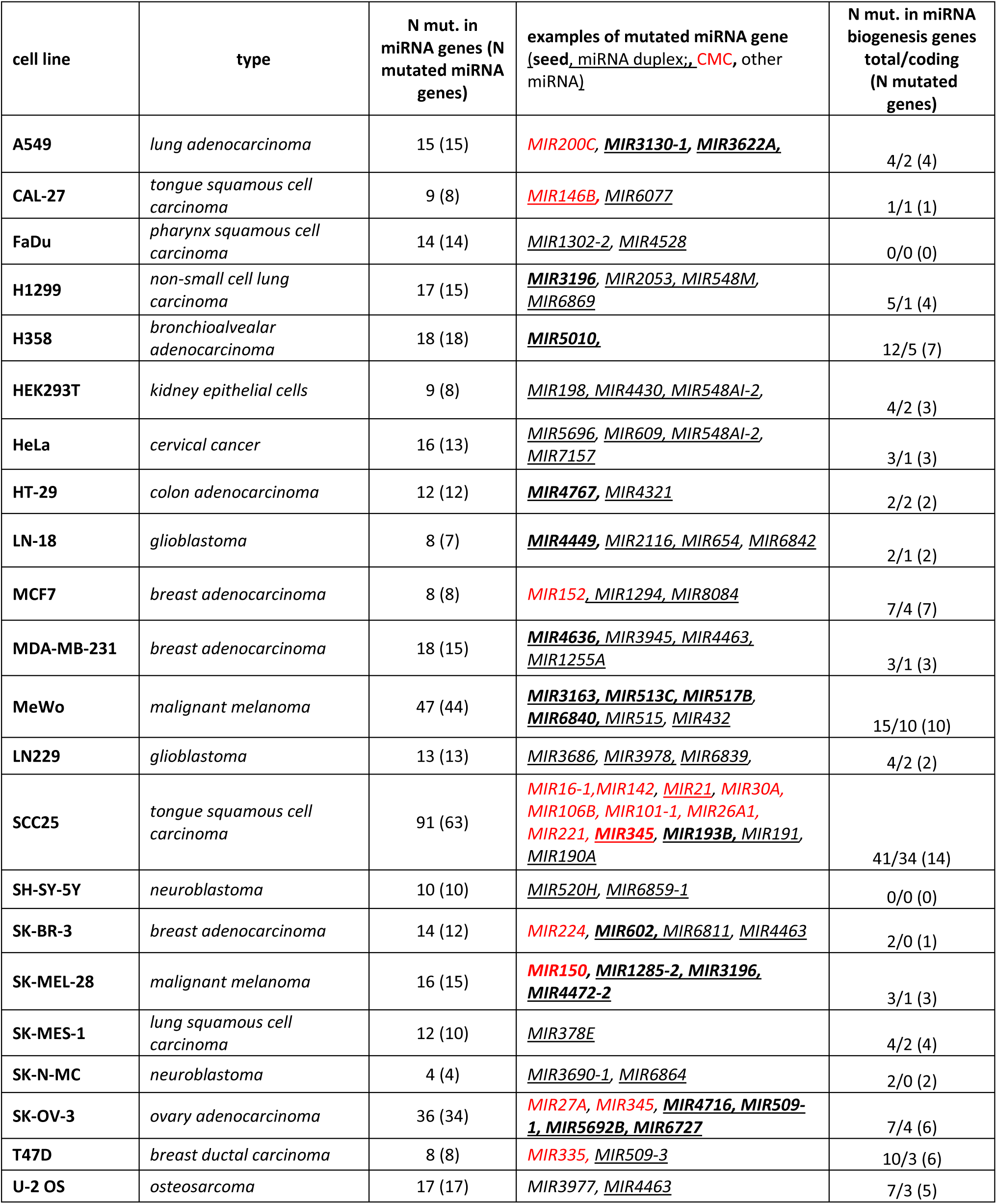
Summary of mutations found in miRNA and miRNA biogenesis genes identified in the selected cancer cell lines.

## DISCUSSION

The WMS developed in this study is the first sequencing system enabling parallel sequencing of all miRNA genes. To demonstrate the potential of WMS, we sequenced several hundred cancer samples of various cancer types and a panel of established cancer cell lines. As a result, we identified nearly 1, 500 somatic mutations in cancer samples and over 500 constitutive mutations in the established cancer cell lines. About two-thirds of these mutations were found in miRNA genes, with the remaining in miRNA biogenesis genes. At least one miRNA gene mutation occurred in 67% of cancer samples. However, this percentage varied between cancer types, with the highest occurring in BCC and LUN and the lowest in REN and OVA, consistent with the general mutational burden in these cancer types (50). Although proportional across cancer types, the fractions of samples with mutations in miRNA genes were somewhat higher in this study than in TCGA samples (14). This discrepancy is likely due to the much higher sequencing depth in this study and the fact that the sequencing of TCGA samples did not cover all miRNA genes The reliability of the detected mutations was verified through various methods, including analyzing mutations using alternative molecular methods and comparing them to data previously published by large-scale and well-recognized projects such as TCGA. All of these analyses showed nearly perfect agreement, or even superiority of WMS over other results, thus confirming the robustness of WMS in detecting mutations with various VAFs in samples of different types, including low-quality archival FFPE samples.

The analysis of identified mutations showed that miRNA gene mutations are evenly distributed along the gene sequence and occur in all miRNA precursor subregions, including flanking sequences, miRNA duplex, and apical loop. Some mutations were identified in miRNA genes’ most crucial functional elements, such as seed sequence, DROSHA and DICER1 cleavage sites or binding motifs of the regulatory RNA-binding proteins. Previous functional analysis of phenotypically neutral single nucleotide polymorphisms (SNPs) located in miRNA genes showed that the majority of sequence variants in miRNA genes affect the accuracy and effectiveness of miRNA processing (19). Therefore, it is justified to expect that a substantial number of mutations identified in this study also have the potential to affect the functioning of miRNA genes. Although few in number, examples of mutations in miRNA genes associated with human diseases, including monogenic Mendelian diseases, demonstrate that sequence variants can impact miRNA genes by disrupting various aspects of their functioning, including the efficiency of miRNA processing/miRNA level, 5p/3p strand ratio, the shift of DROSHA/DICER1 cleavage sites generating alternative isomiRs, and the efficiency of recognition and silencing of the target genes (summarized and discussed (15)). It is anticipated that such effects can be obtained either by the direct impact of mutations changing the key functional sequences of miRNA genes, such as seed or protein-binding sites or by the influence of mutations on the structure and stability of miRNA precursors. Nonetheless, it has to be noted that since most mutations in the cancer genome are randomly occurring neutral variants, only a small fraction of mutations detected in miRNA genes may play a role in carcinogenesis, regardless of their impact on miRNA genes. The most potent in this term may be mutations in well-known cancer-related miRNA genes, such as *MIR16-2*, *MIR143,* or *MIR155*, or mutations recurring in specific miRNA genes.

Although the analysis of miRNA gene mutations with OncodriveFML had low statistical power, it identified several miRNA genes with an enriched signal of mutation functionality, suggesting their potential as cancer drivers, with *MIR3928* showing the most substantial evidence. The mutations in *MIR3928* occur specifically in BCC, with 4 mutations in 4 different samples. All mutations are located in the 5p-flanking region, including one singleton mutation and one hotspot mutation (occurring in 3 samples) located, respectively, 21 and 18 nucleotides upstream of the pre-miR-3928 sequence. Computational modeling showed that the mutations, especially the hotspot mutation, locally modify the structure and decrease the stability of the miR-3928 precursor. *MIR3928* is a well-validated miRNA gene (miRBase); however, it has not been strongly implicated in cancer, except for one recent study suggesting the role of miR-3938 in glioma (51). However, it must be noted that *MIR3928* should only be considered a candidate cancer driver, and the oncogenic nature of its mutation should be interpreted cautiously. Proving the oncogenic/driver nature of mutations in a gene would require additional experimental and clinical analyses as well as testing the gene in additional larger panels of samples, which are beyond the scope of our study.

In comparison, the strongest candidate cancer driver identified among the miRNA biogenesis genes is *SMAD4*. This well-known tumor suppressor plays a role and is commonly mutated in various solid tumors, particularly in pancreatic and other intestinal duct cancers (22, 33, 34). The identification of *SMAD4* confirms the reliability of the collected data and the performed analysis. SMAD4 is a well-characterized transcription factor that functions as a heterotrimer formed with other SMAD family members and regulates the transcription of various genes, including miRNA genes (52, 53). Mutations in *SMAD4*, especially in the MH2 domain, prevent the interaction of SMAD4 with other SMADs and, thus, the formation of the functional complex (22).

Like WES, WMS can identify various types of mutations in different types of human genomic samples, including germline and somatic mutations, whether rare or common, and with different VAF. Particularely the WMS data may also be used to identify large-scale CNAs. Particularly, WMS may be used to identify mutations in diseases or conditions with “hidden heritability, “ in which mutations in coding genes and other types of genetic variation routinely tested for diseases have been excluded. Such cases may include mutations in *MIR96* (11, 12) and *MIR204* (54) identified in patients with hereditary conditions, nonsyndromic hearing loss, and retinal dystrophy in which no mutations in known genes associated with the conditions were identified. Another application of WMS may be the in-depth characterization or screening of mutations in a miRNA gene or group of genes in which mutations were already detected or suspected. Such genes may include *MIR14*2, with somatic mutations in various blood cancers (14), or *MIR15A and MIR16-1,* commonly deleted and/or mutated in chronic lymphocytic leukemia (13). WMS may also be used to identify new genes responsible for cancer predisposition. For example, although many genes have been already identified and are being tested for breast cancer predisposition, new genes are still being sought to explain all the suspected heritable cases. So far, all these analyses have ignored noncoding genes, including miRNA genes. Next, WMS may be used to prescreen samples or cell lines before selecting them for the functional study of miRNA or miRNA-related processes to avoid using a model with a mutation in the miRNA of interest. Finally, WMS may be used as a complementary test to miRNA expression profiling, as, on the one hand, miRNA gene mutations may affect miRNA levels, and on the other hand, miRNAs with unchanged levels may not be functional due to mutations, e.g., in seed sequence.

The additional value of our study is the catalog of miRNA gene mutations in established cancer cell lines. Some of these mutations are located in extensively studied cancer-related genes (e.g., *MIR16-1* and *MIR21* in SCC25, *MIR152* in MCF7, *MIR150* in SK-MEL-28, or *MIR345* in SK-OV-3). This information may be essential and should be considered when selecting a cell line for a miRNA functional study or interpreting the functional results in a particular cell line. Similarly, the mutations in miRNA biogenesis genes (e.g., nonsense mutations in *DICER1* in H358 and *SMAD4* in CAL27 and HT-29) may be considered when performing relevant experiments.

The primary benefit of using WMS is the much lower cost of sequencing miRNA genes in the human genome than the only available alternative, WGS. For example, the cost of our experiment of sequencing 580 samples with ∼700x coverage, including library preparation and sequencing, equaled 133, 400$ (230$ per sample). The cost may be further substantially reduced by using newer machines and applying higher indexing/multiplexing of samples. It allows for sequencing with much higher coverage or analysis of a much larger number of samples. An additional advantage of the WMS system is the relatively small amount of DNA sample required. In the study, we used 150-200 ng for sequencing, while the minimum amount recommended for WGS is about ten times higher (2 µg). The low sample consumption substantially increases the applicability of WMS for low-quantity samples, which are often cancer samples but also other human specimens. Finally, in our experiment, we demonstrated the applicability of WMS for different types of DNA samples, including very low-quality, highly degraded FFPE cancer samples.

In conclusion, the study involved the development of WMS, a tool for targeted sequencing of all human miRNA genes. We used WMS to sequence over 300 cancer samples, as well as cell line samples and identified 1, 291 mutations in miRNA genes. The high reliability of the detected mutations and the robustness of WMS were confirmed by various methods. However, it is important to note that interpreting the functional consequences of sequence variants in noncoding sequences, such as miRNA genes, remains challenging. Improving this area will require collecting more data, developing new computational and statistical approaches, and expanding genetic knowledge. Tools like WMS can assist in gathering relevant data, ultimately contributing to understanding the consequences of mutations in miRNA genes.

## DATA AVAILABILITY

All data generated or analyzed during this study are included in the published paper and associated supplemental files. WMS is available from Agilent SureSelect Custom target capture under design ID:3115731 upon request from the corresponding author.

## AUTHORS’ CONTRIBUTIONS

Paulina Galka-Marciniak: Conceptualization, Data curation, Formal analysis, Methodology, Resources, Validation, Visualization, Writing – original draft, Writing – review & editing. Martyna O. Urbanek-Trzeciak: Resources, Software, Methodology, Writing – review & editing. Daniel Kuznicki: Formal analysis, Software, Resources, Writing – review & editing. Natalia Szostak: Resources, Software, Writing – review & editing. Adrian Tire: Validation, Investigation, Visualization, Writing – review & editing. Paulina M. Nawrocka-Muszynska: Investigation, Validation, Writing – review & editing. Katarzyna Chojnacka: Resources, Writing – review & editing. Malwina Suszynska: Conceptualization, Methodology, Writing – review & editing. Katarzyna Klonowska: Conceptualization, Writing – review & editing. Karol Czubak: Conceptualization, Writing – review & editing. Magdalena Machowska: Conceptualization, Writing – review & editing. Anna Philips: Resources, Writing – review & editing. Konstantin Maksin: Resource, Writing – review & editing. Laura Susok: Resources, Writing – review & editing. Michael Sand: Resources, Writing – review & editing. Janusz Rys: Resources, Writing – review & editing. Jolanta Jura: Resources, Writing – review & editing. Magdalena Ratajska: Resources, Writing – review & editing. Hanna Dams-Kozlowska: Resources, Writing – review & editing. Janusz Kowalewski: Resources, Writing – review & editing. Marzena Anna Lewandowska: Resources, Validation, Writing – review & editing. Piotr Kozlowski: Conceptualization, Project administration, Supervision, Resources, Funding acquisition, Formal analysis, Methodology, Writing – original draft, Writing – review & editing.

## ACKNOWLEDGEMENTS

We would like to acknowledge the Poznan Supercomputing and Networking Center (PSNC) for the computing resources (Eagle supercomputer) supporting this work.

## SUPPLEMENTARY FIGURES

**Supplementary Figure S1.**
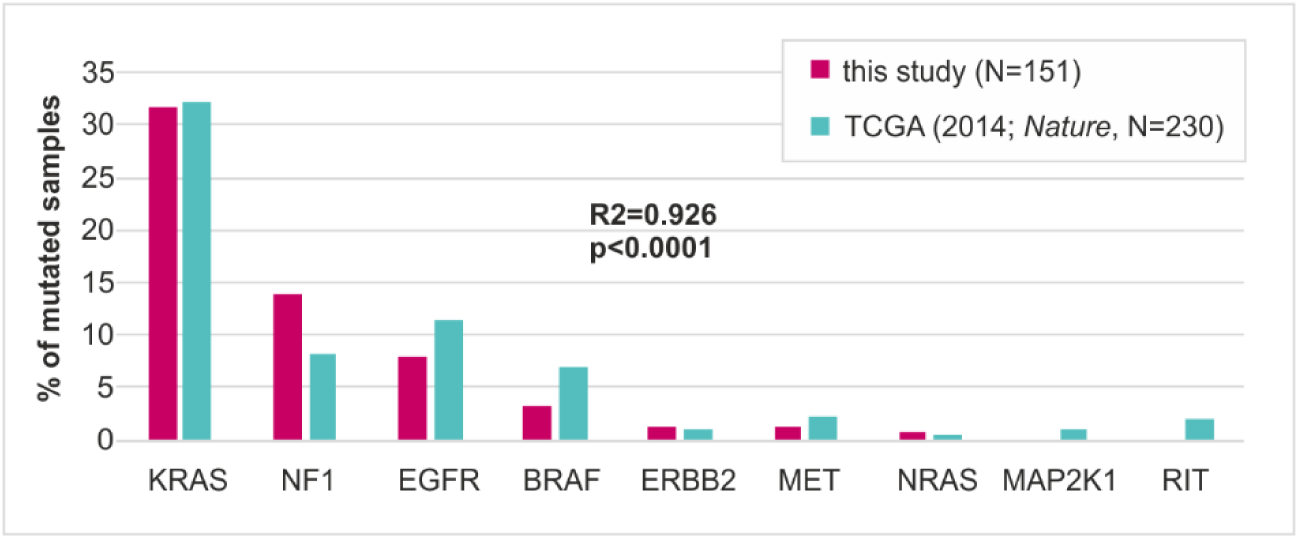
Comparison of mutation frequency in lung adenocarcinoma driver genes identified in this study in LUN samples (n=151) and TCGA lung adenocarcinoma project (n=230) (2014, Nature). The Pearson correlation coefficient (R2) and p-value are indicated on the graph.

**Supplementary Figure S2.**
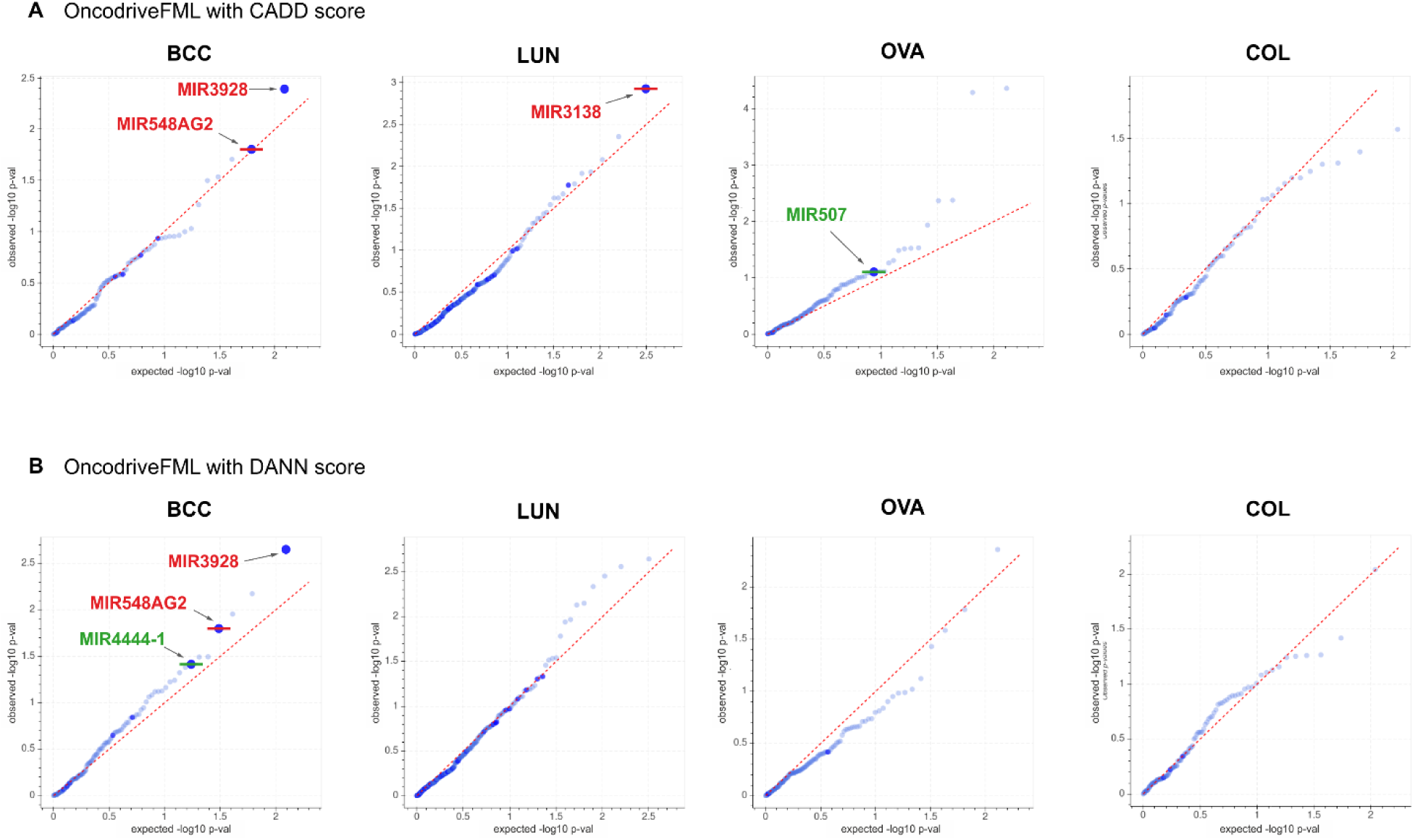
Analysis of cancer driver potential in miRNA genes performed with OncodriveFML separately for mutations identified in BCC, LUN, OVA, and COL. The QQ plots show the distribution of expected (x-axis) and observed (y-axis) p-values corresponding to functional mutation bias calculated with (A) CADD and (B) DANN scores. The green and red colors indicate genes defined as significant (q<0.25) and highly significant (q<0.1), respectively, according to the OncodriveFML recommendation.

**Supplementary Figure S3.**
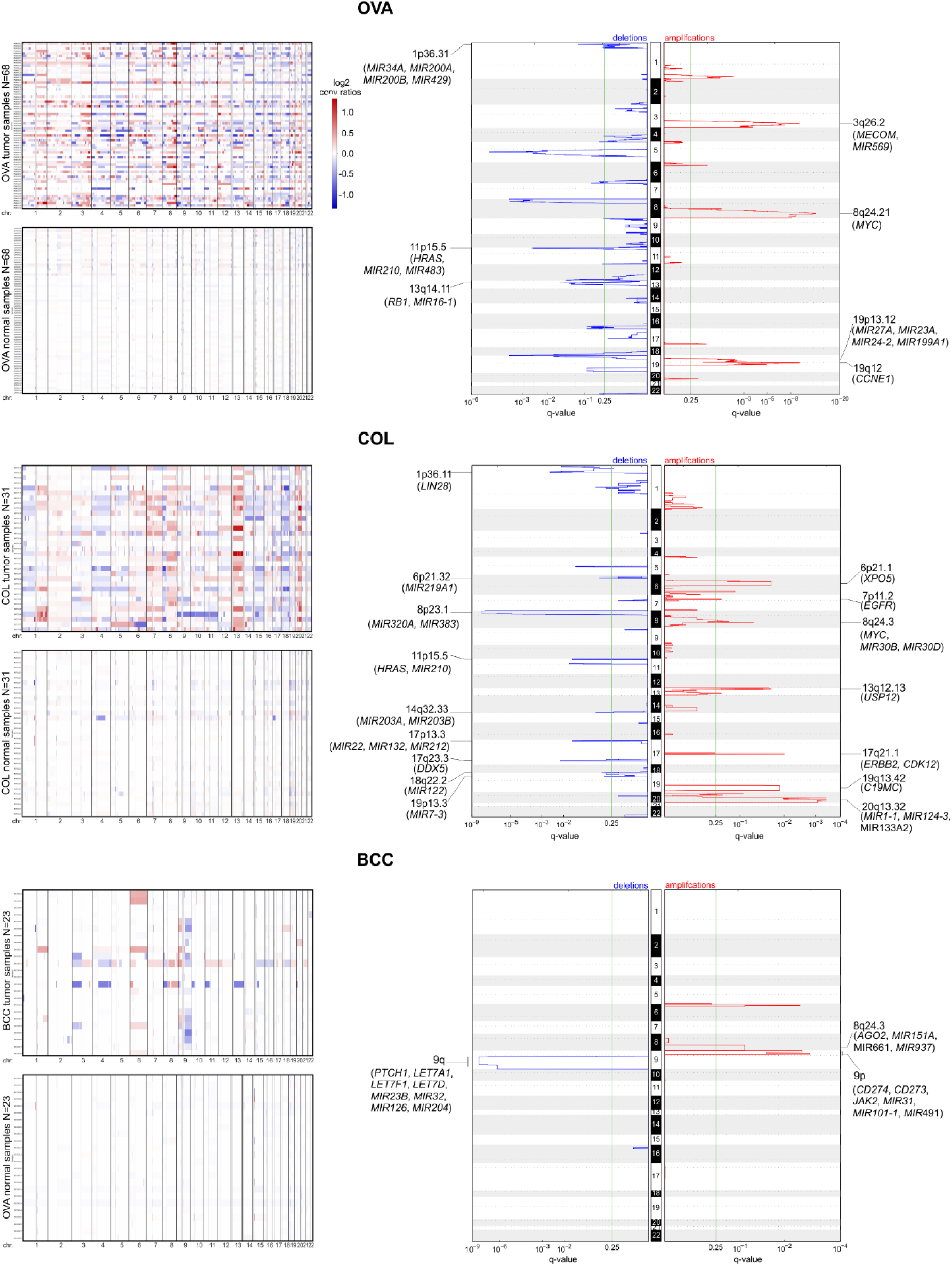
CNA analysis using WMS data in OVA, COL, and BCC samples. The figure scheme as in Figure 6.

